# Proteomic Signatures of Neuropathological Alterations in Alzheimer’s Disease: Insights from the Bio-Hermes Study

**DOI:** 10.64898/2026.05.20.726478

**Authors:** Hanjun Zhao, Taiyu Zhu, Buddhiprabha Erabadda, Ganna Leonenko, Rashmi Maurya, Dicky Lim, Kalliopi Mavromati, Terry Quinn, Ivan Koychev, Valentina Escott-Price, Shisong Jiang, Alejo Nevado-Holgado, Laura Winchester

## Abstract

Large-scale plasma proteomics can capture molecular changes across the Alzheimer’s disease (AD) continuum and provide insight into biological mechanisms associated with AD pathology. We analysed the Bio-Hermes cohort (n = 961), with participants enrolled across 17 sites in the United States from April 2021 to November 2022. Participants were stratified by clinical status and amyloid PET scan-based Core1 biomarker status (CN Core1**–**, CN Core1**+**, MCI Core1**+**, and AD dementia Core1**+**). We performed differential abundance analyses across biologically defined contrasts, clustered proteins into co-expression networks, and evaluated protein panels to distinguish participants with biologically defined AD from amyloid-negative cognitively normal controls. We also used Mendelian randomization (MR) to assess genetic evidence for potential causal relationships with AD risk. The biologically defined contrast, Core1**+** vs. CN Core1**–**, identified 69 differentially abundant proteins. Across AD stages, eight core proteins were consistently dysregulated from preclinical through prodromal and dementia phases, and three additional proteins emerged at MCI Core1**+** and remained altered in AD dementia Core1**+**. We identified 29 co-expression modules, six of which varied significantly across the AD continuum. Among differential abundance proteins, ACHE ranked highest for distinguishing biologically defined AD from CN Core1**–**. Stage-specific protein panels improved the discriminatory performance for MCI Core1**+** (AUC = 0.850) and AD dementia Core1**+** (AUC = 0.856). MR provided genetic evidence consistent with an association between plasma ACHE abundance and AD risk. Plasma proteomics delineated a stage-spanning core signature across the AD continuum. These findings nominate co-expression modules and candidate proteins for further validation in early detection and AD screening.

## 1 Introduction

Alzheimer’s disease (AD) is a progressive neurodegenerative disorder that begins with the emergence of AD neuropathologic change (ADNPC) [1]. The pathological cascade is initiated by the accumulation of extracellular deposits of amyloid-*β* (A*β*) plaques and followed by the aggregation of intracellular misfolded tau proteins [2]. Despite the clinical utility of these hallmarks, they capture only a portion of the complex pathophysiological processes of AD [3]. Moreover, the characteristic neuropathological changes of AD can arise two decades or more before the onset of clinical symptoms, such as memory loss and cognitive decline [4]. Currently, our understanding of the pathological alterations and the molecular transition from preclinical AD and prodromal AD to dementia due to AD remains limited [5]. Therefore, comprehensive proteomic profiling across the AD continuum may deepen insight into pathological mechanisms and support the discovery of biomarkers and candidate therapeutic targets to delay progression.

Amyloid positron emission tomography (PET) provides an accurate measurement of amyloid pathology (moderate to frequent neuritic A*β* plaques) validated against autopsy. It aligns with criteria for intermediate/high ADNPC [6] and is used extensively in AD clinical research and drug development [7]. Using radiotracers with high affinity for fibrillar amyloid aggregates [8], amyloid PET is a non-invasive modality and has been designated a Core1 biomarker for biologically defined AD diagnosis in the updated version of the National Institute of Aging-Alzheimer’s Association (NIA-AA) criteria [1].

Recent advancements in proteomics have enabled the simultaneous quantification of a wide range of proteins [9]. Unbiased high-throughput proteomic profiling provides a data-driven framework for identifying peripheral molecular alterations implicated in AD and prioritising associations with clinically and neuropathologically relevant traits [10]. Despite rapid growth in AD proteomic research [11, 12], studies that explicitly anchor plasma proteomic signatures to disease-specific Core biomarkers supporting the biological definition of AD remain limited.

The Bio-Hermes Study is a well-characterised large-scale cohort comprising cognitively normal (CN), mild cognitive impairment (MCI) and AD dementia participants [13, 14]. Participants provided peripheral blood samples for genetic and proteomic analyses and underwent amyloid PET imaging and clinical assessments, including cognitive evaluations and plasma brain-derived biomarkers, yielding a multimodal dataset with rich endophenotypic traits [13]. By integrating proteomics with amyloid PET, Bio-Hermes offers a framework to delineate stage-specific proteomic signatures throughout the AD continuum. This approach may provide insights into mechanisms underlying pathological progression, reduce biological heterogeneity and support discovery of candidate molecules for prognostic biomarker development and mechanistic follow-up.

We integrated plasma proteomic profiling using SomaScan with amyloid PET imaging to quantify the presence and magnitude of AD pathology. We pursued four objectives: (1) to identify proteins that differ significantly in biologically defined AD [1, 9] and across AD continuum stages; (2) to conduct network and pathway enrichment analyses to define the proteomic patterns, highlighting shared functional relationships and associations with AD endophenotypes; (3) to construct and validate the protein-based diagnostic models for distinguishing biologically defined AD from controls; (4) to perform Mendelian randomization (MR) analyses to evaluate genetic evidence for potential causal relationships between dysregulated protein abundance and AD risk.

## 2 Results

### 2.1 Participants’ characteristics

Plasma proteomics and AD endophenotypes, including cognitive assessments and brain-derived biomarkers, were analysed in 961 participants from the Bio-Hermes Study. Based on the 2024 Revised Criteria of the Alzheimer’s Association Workgroup [1], participants were stratified by clinical status and Core1 biomarker status, assessed using amyloid PET. Six groups were defined: CN Core1– (n = 306), CN Core1+ (n = 81), MCI Core1– (n = 183), MCI Core1+ (n = 104), non-AD dementia Core1–(n = 95), and AD dementia Core1+ (n = 141). Amyloid PET data were unavailable for 51 participants. Primary analyses focused on biologically defined AD, comprising all participants classified as CN Core1+, MCI Core1+, or AD dementia Core1+ [1, 9]. Subsequently, distinct stages of the AD continuum, from CN Core1+ and MCI Core1+ to AD dementia Core1+, were investigated. Figure 1 provides an overview of the study design and main analyses.

**Fig. 1.**
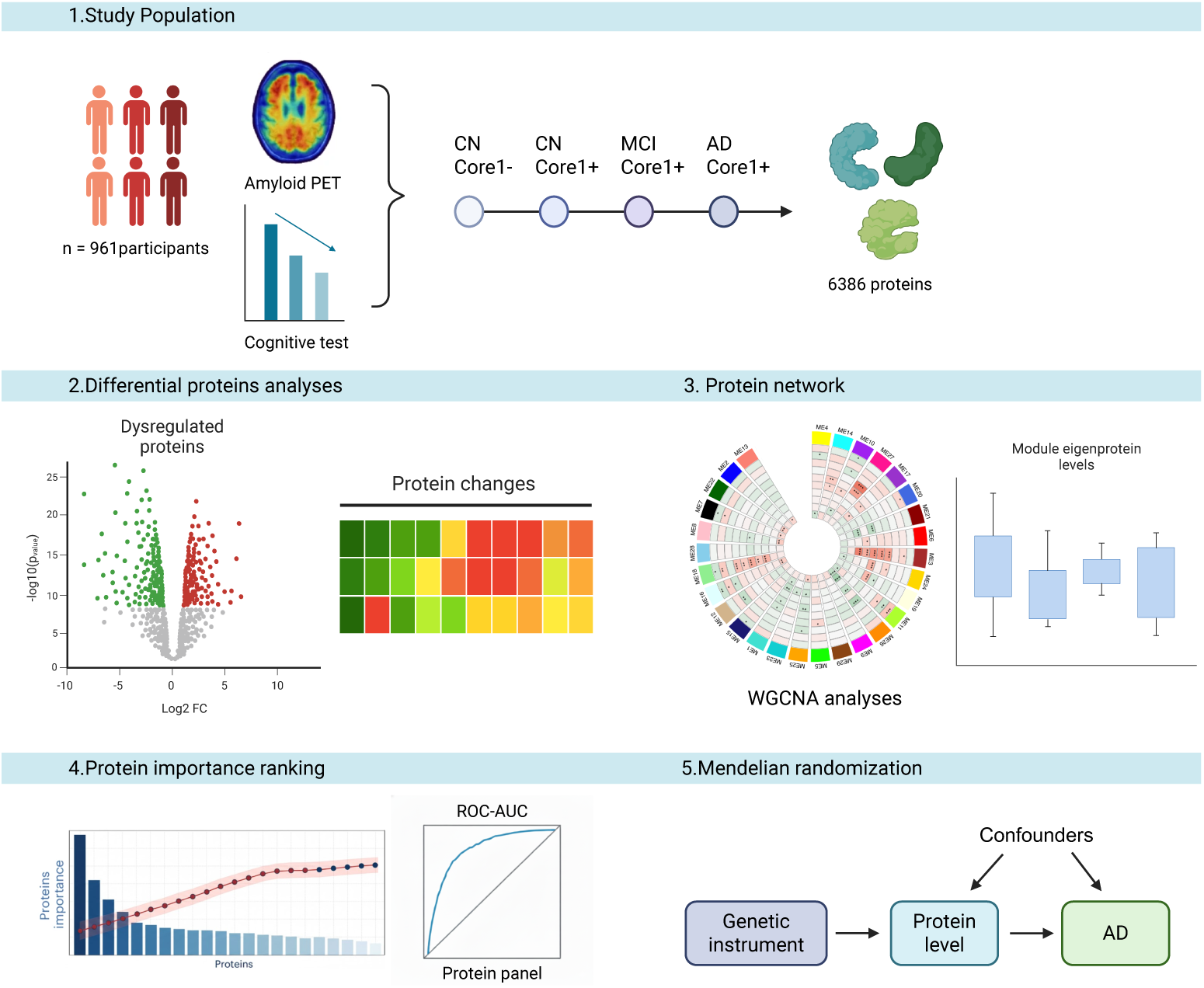
Study overview. From the Bio-Hermes Study, we included 961 participants with proteomics data. (1) We performed differential abundance analyses on 6,386 proteins between distinct stages of biologically defined AD and CN Core1–, identifying proteins that differed across the AD continuum; (2) we conducted network and pathway enrichment analyses to define proteomic patterns, shared functional relationships, and associations with AD endophenotypes; (3) we developed protein-based classification models for distinguishing biologically defined AD from controls and assessed the discriminatory performance of the proteomic panels; (4) we performed Mendelian randomization (MR) analyses to assess the genetic evidence for potential causal relationships between dysregulated proteins and AD risk. WGCNA, weighted gene co-expression network analysis; ROC, receiver operating characteristic.

The demographic characteristics of participants stratified by clinical status and amyloid PET modality are summarised in Table 1. No significant difference in sex distribution was observed among groups. By contrast, groups across the AD continuum differed in age, education, and *APOE-*ε*4* carrier prevalence. Cognitive assessments also differed across biologically defined groups, with MMSE scores declining and FAQ scores increasing across disease stages. Detailed demographics for the clinical diagnostic groups in the Bio-Hermes Study are provided in Supplementary Table 2.

**Table 1.** Demographic characteristics stratified by Core1 biomarker status and clinical diagnosis.

| Characteristics | CN |  | MCI |  | Non-AD | AD dementia | P value |
| --- | --- | --- | --- | --- | --- | --- | --- |
|  | Core1−<br>(n=306) | Core1+<br>(n=81) | Core1−<br>(n=183) | Core1+<br>(n=104) | Core1−<br>(n=95) | Core1+<br>(n=141) |  |
| Age, years | 69 (65–74) | 73 (67–78) | 70 (65–77) | 75 (70–78.25) | 74 (70–78) | 75 (72–79) | <0.001 |
| Sex (male) | 193 (63.07%) | 46 (56.79%) | 101 (55.19%) | 55 (52.88%) | 54 (56.84%) | 72 (51.06%) | 0.181 |
| Education, years | 16 (14–18) | 16 (14–17) | 16 (14–17) | 16 (13–18) | 15 (13–17) | 15 (12–18) | 0.020 |
| RAVLT | 45 (38–51) | 40 (35–47) | 35 (30–40) | 32 (27–38) | 28 (24–36) | 24 (19–29) | <0.001 |
| MMSE | 29 (27–30) | 29 (28–29) | 28 (26–29) | 27 (25–29) | 24 (22–25) | 23 (21–25) | <0.001 |
| FAQ | 0 (0–1) | 0 (0–2) | 2 (0–4) | 4 (1–6) | 7 (3–12) | 9 (5–13) | <0.001 |
| APOE-ε4 carriers | 79 (25.82%) | 48 (59.26%) | 43 (23.5%) | 67 (64.42%) | 12 (12.63%) | 91 (64.54%) | <0.001 |

Note: Continuous variables are presented as median (interquartile range), and categorical variables are presented as n (%). P values for age, education, MMSE, RAVLT, and FAQ were generated using Kruskal–Wallis rank-sum tests; P values for sex and APOE-ε4 carrier status were generated using Pearson’s chi-square tests.
APOE, apolipoprotein E; FAQ, Functional Activities Questionnaire; MMSE, Mini-Mental State Examination; RAVLT, Rey Auditory Verbal Learning Test.

### 2.2 Plasma proteomic differences in AD

Plasma proteomic profiling in the Bio-Hermes Study was performed by the SomaScan 7K panel (SomaScan v4.1 assay). After quality control, 7,289 aptamers mapping to 6,386 unique human proteins were retained. To identify circulatory proteins associated with clinically diagnosed AD dementia, we fitted multiple linear regression models adjusted for age, sex, and the first two proteomic principal components (PCs) to account for potential experimental confounding factors (model 1; four-covariate model) [11]. We identified 451 proteins (468 aptamers) that were differentially abundant in clinically diagnosed AD dementia group relative to cognitively normal controls (Benjamini-Hochberg false discovery rate (FDR)-adjusted *P* value; *P*_FDR_ *<* 0.05), comprising 192 upregulated proteins (198 aptamers, 42.6%) and 259 downregulated proteins (270 aptamers, 57.4%) (Fig. 2a, Supplementary Table 4). Acetylcholinesterase (ACHE) was the top-ranked upregulated differentially abundant protein (DAP), followed by VAV3 and TEC. The leading downregulated DAPs were RAB5B, ATE1, and SUGT1.

**Fig. 2.**
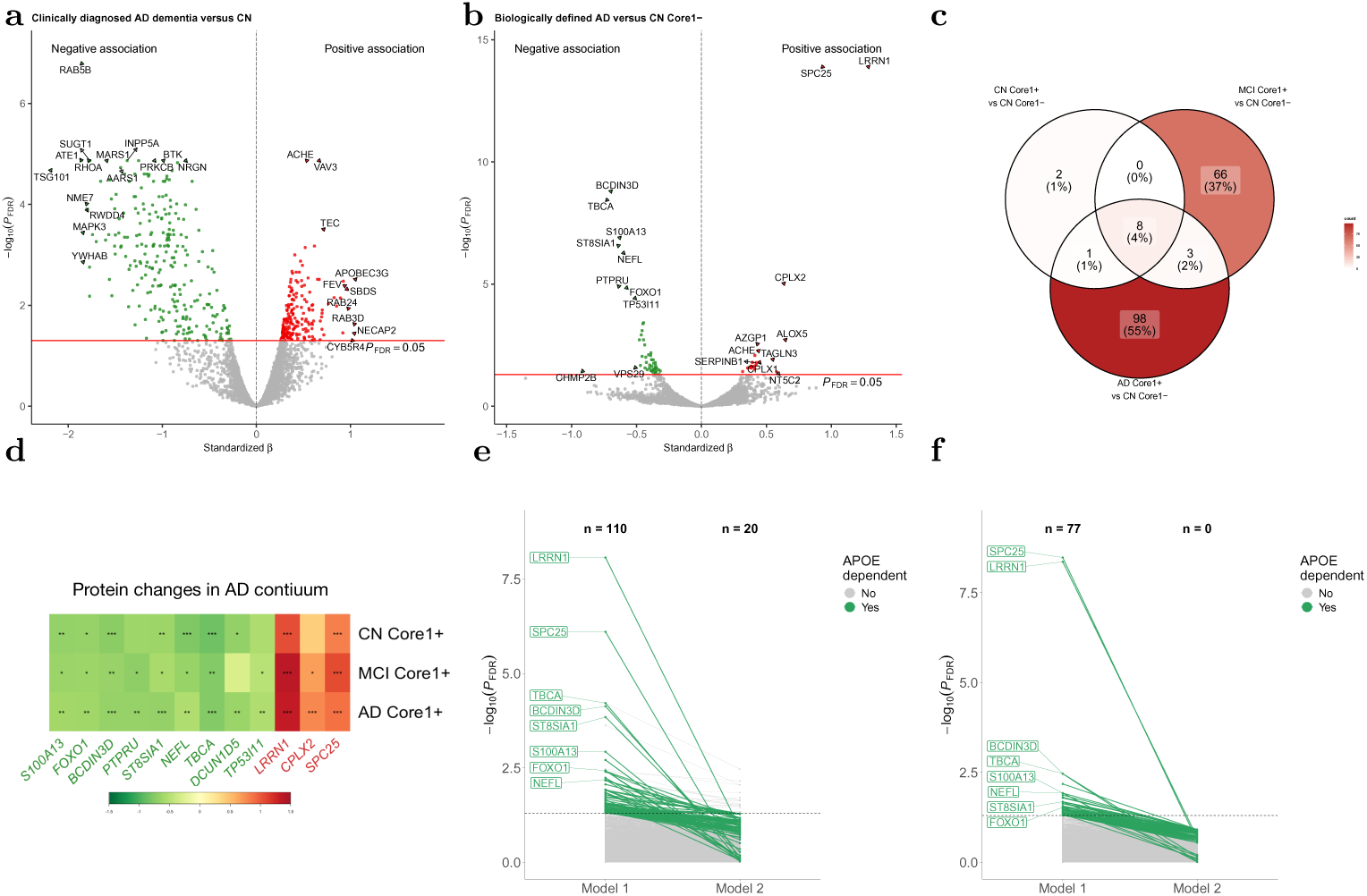
Differentially abundant proteins (DAPs) across the AD continuum. **a,b,** Volcano plots showing standardised *β* (x axis) and *−* log_10_ *P*_FDR_ (y axis) for the differential analyses comparing clinically diagnosed AD dementia with CN (**a**) and biologically defined AD with CN Core1– (**b**). Multiple linear regression models included age, sex, and the first two proteomics PCs as covariates. The red horizontal line indicates the threshold of *P*_FDR_ *<* 0.05. **c,** Venn diagram showing overlap number in significant dysregulated proteins from CN Core1+, MCI Core1+, and AD dementia Core1+. **d,** Heat map showing changes in overlapping dysregulated proteins across the AD continuum. Colours represent standardised *β* and asterisks indicate statistical significance after Benjamini–Hochberg FDR correction (^no symbol^*P ≥* 0.05, * *P <* 0.05, ** *P <* 0.01, *** *P <* 0.001). **e,f,** Spaghetti plots showing Benjamini–Hochberg FDR-adjusted significance for protein associations with AD dementia Core1+ (**e**) and MCI Core1+ (**f**) across the two linear regression models, highlighting eight overlapping proteins (green) whose associations with distinct AD stages were attenuated upon *APOE-*ε*4* adjustment.

To investigate plasma proteins associated with biologically defined AD, differential abundance analyses were conducted following the same protein selection procedures. We identified 69 proteins (71 aptamers) significantly associated with biologically defined AD (*P*_FDR_ *<* 0.05; Fig. 2b, Supplementary Table 5). The top DAPs were LRRN1, SPC25, BCDIN3D, TBCA, S100A13, ST8SIA1, NEFL and CPLX2. Among these, LRRN1, SPC25, NEFL, TBCA, and S100A13 recapitulated findings from recent AD proteomics studies [11, 12, 15, 16], whereas BCDIN3D, ST8SIA1 and CPLX2 emerged as novel candidates from the Bio-Hermes Study. To evaluate proteins associated with both clinically diagnosed AD dementia and biologically defined AD, we identified 11 proteins significant in both analyses, each with directionally concordant associations (Extended Data Fig. 2a-2b, Supplementary Tables 4-5).

In sex-stratified subgroup analyses of clinically diagnosed AD dementia, we identified 143 DAPs (149 aptamers) in females and two in males (*P*_FDR_ *<* 0.05; Supplementary Table 6). In biologically defined AD, 26 DAPs (27 aptamers) were observed in females and four in males. Four proteins (including LRRN1, SPC25, TBCA, and S100A13) were robustly dysregulated across sex strata and remained concordant with primary findings in all Core1+ groups (Supplementary Table 7), suggesting that circulatory protein profiles were broadly comparable across sex groups. Within the female group, the findings showed both substantial overlap with and divergence from the entire population, whereas C14orf93, TMCC3, and KRT19 were significantly dysregulated only among women.

### 2.3 Plasma protein changes across AD stages

To investigate protein alterations across the pathological cascade of AD, we analysed plasma DAPs at each disease stage. Eight overlapping proteins (LRRN1, SPC25, BCDIN3D, TBCA, S100A13, ST8SIA1, NEFL, and FOXO1) were consistently dysregulated across all three stage-wise comparisons (Fig. 2c-2d, Extended Data Fig. 2c-2d). Two proteins (PPP1R8 and ITGA4/ITGB1) and 66 proteins (including SKIL, TAC4, AZGP1, and VLDLR) were uniquely differentially abundant in the CN Core1+ and MCI Core1+ phases versus CN Core1–, respectively. These findings suggest that some stage-specific effects in preclinical and prodromal AD may not persist as disease progresses to dementia due to AD. Three proteins (CPLX2, PTPRU and TP53I11) became significant in the MCI Core1+ phase and remained altered during the AD dementia Core1+ phase, highlighting characteristic changes emerging in later stages of the disease. Most DAPs (*n* = 98) were uniquely associated with AD dementia Core1+ group, demonstrating substantial proteomic remodelling in advanced disease (Extended Data Fig. 3b-3d). Summary statistics from the differential analyses were reported in Supplementary Table 8.

**Fig. 3.**
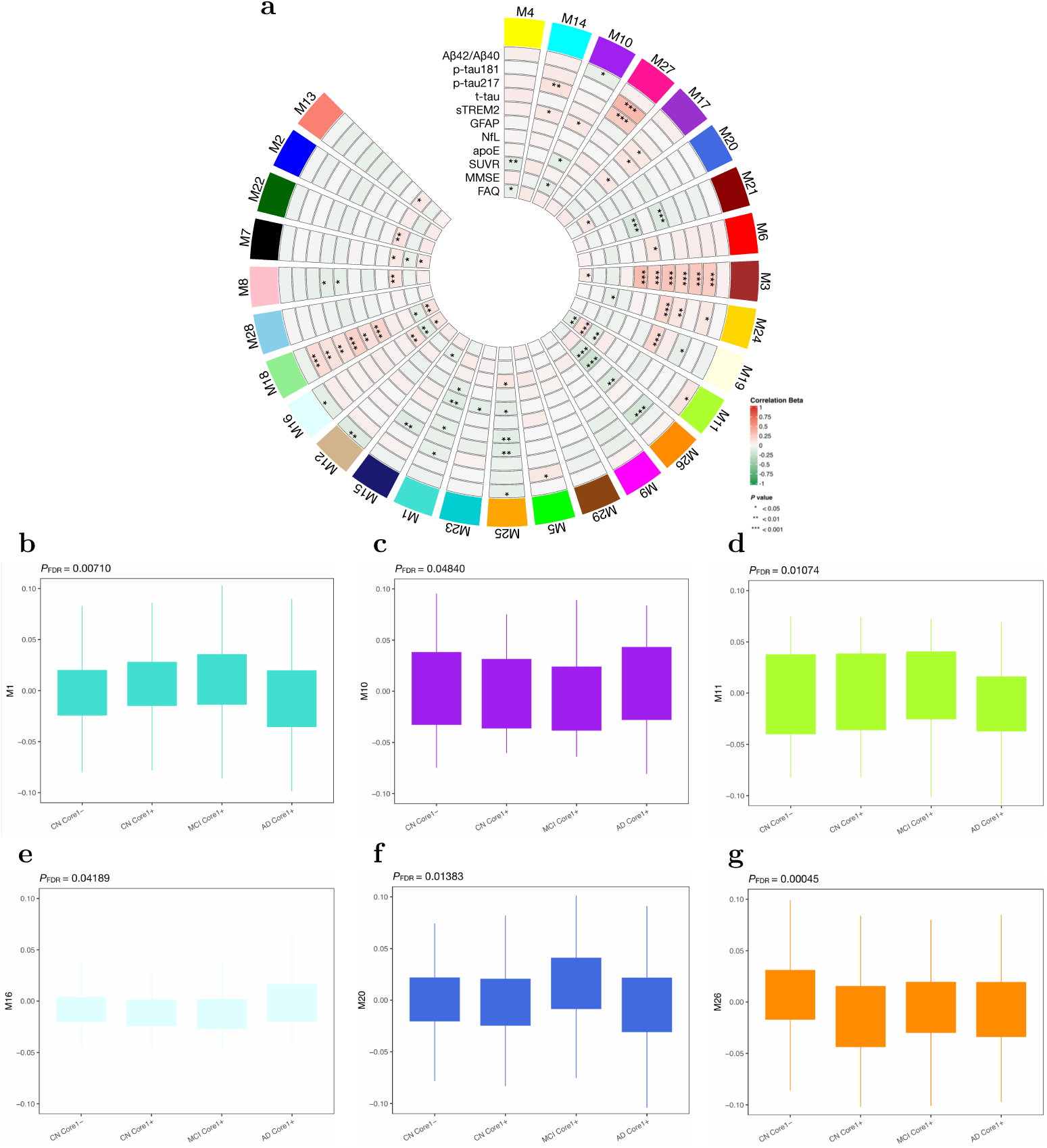
AD protein co-expression network reveals modules linked to AD endophenotypes and stages. **a,** A protein co-expression network was built using WGCNA and comprised 29 co-expression modules. Module eigenproteins were correlated with blood-based neuropathological, cognitive, and imaging traits present in the Bio-Hermes Study (red indicates positive correlation; blue indicates negative correlation). Among the 29 modules, 26 showed significant correlations with at least one clinical feature. Colour intensities represent Pearson correlation coefficients and asterisks indicate statistical significance after Benjamini–Hochberg FDR correction (^no symbol^*P ≥* 0.05, * *P <* 0.05, ** *P <* 0.01, *** *P <* 0.001) **b-g,** Module eigenprotein levels by AD continuum for the six significantly associated modules. Groups are ordered from CN Core1- and CN Core1+ through MCI Core1+ to AD dementia Core1+. Differences in module eigenprotein levels among groups were evaluated using the Kruskal-Wallis rank-sum tests.

*APOE-*ε*4* genotype is strongly associated with AD risk [17]. To assess the impact of *APOE-*ε*4* on protein alterations, we re-evaluated DAPs with additional adjustment for the *APOE-*ε*4* carrier status in sensitivity analyses (model 2; five-covariate model). We found 20 DAPs significantly associated with the AD dementia Core1+ phase (*P*_FDR_ *<* 0.05, Fig. 2e, Extended Data Fig. 2e-2f, Supplementary Table 9). Comparing models, 17.3% (19 of 110) of model 1 DAPs remained significant after *APOE-*ε*4* adjustment, indicating that their associations with AD dementia Core1+ were independent of *APOE-*ε*4* carrier status. Leading *APOE-*ε*4* -independent proteins included ACHE, RAB1A, CPLX2, SERPINA3, HSD17B7, and VAV3. The remaining 82.7% (91 of 110) DAPs became non-significant after *APOE-*ε*4* adjustment and were classified as *APOE-*ε*4* -dependent proteins, including the eight overlapping DAPs (including LRRN1, SPC25, BCDIN3D, TBCA, S100A13, ST8SIA1, NEFL, and FOXO1). One protein, AP1S2, gained significance after adjustment. In the MCI Core1+ group, all DAPs became non-significant after *APOE-*ε*4* adjustment (*P*_FDR_ *<* 0.05; Fig. 2f, Supplementary Table 9), suggesting that a large proportion of AD-associated DAPs across disease stages were influenced by the *APOE-*ε*4* genotype.

To gain insight into cell-type specificity, we quantified the proportion of messenger RNA (mRNA) expression of DAP-encoding genes across major brain cell types using single-cell transcriptomics data from the Allen Human Brain Atlas (AHBA) [18]. Most DAPs were predominantly expressed in neurons, distributed across glutamatergic excitatory and *γ*-aminobutyric acid (GABA)-ergic inhibitory populations, regardless of AD stage. A subset of DAPs showed distinct expression profiles in non-neuronal cell types. Among the 13 overlapping proteins dysregulated in at least two stages, LRRN1 was enriched in oligodendrocyte precursor cells (OPCs), whereas FOXO1 was enriched in astrocytes. SMOC1 and PIP4K2A, uniquely dysregulated in AD dementia Core1+, were specific to OPCs and oligodendrocytes, respectively. PIP4K2A and VAV3, both upregulated in AD dementia Core1+, as well as PRKG1, upregulated in MCI Core1+, were predominantly expressed in astrocytes (Extended Data Fig. 3a-3e).

### 2.4 AD protein co-expression network

To characterise the multifaceted pathophysiology of AD and delineate underlying biological pathways, we constructed a protein co-expression network using the Weighted Gene Co-expression Network Analysis (WGCNA) algorithm. We identified 29 co-expressed protein modules with similar expression patterns, ranked by size from the largest (M1; n = 1098 proteins) to the smallest (M29; n = 15 proteins) (Fig.3a, Supplementary Tables 10-11). These modules were independently recapitulated using *t* -distributed stochastic neighbour embedding (t-SNE), indicating that WGCNA-defined protein communities were robust (Extended Data Fig. 4a). We further validated the protein co-expression network using an independent cross-sectional AD cohort from the Global Neurodegeneration Proteomics Consortium (GNPC) version 1.1 Contributor C cohort. The GNPC is the largest harmonized neurodegeneration proteomic dataset, consisting of biofluid samples profiled using the SomaLogic platform for protein quantification [19, 20]. Twenty-eight of 29 WGCNA modules were highly preserved in the target cohort (Extended Data Fig. 4b), highlighting consistency of expression patterns across cohorts.

To assess whether co-expression modules were related to AD endophenotypes, we correlated the module eigenproteins with plasma-based brain-derived (PBBD) biomarkers, amyloid PET standardised uptake value ratios (SUVR), and cognitive assessments. The investigated clinical features included A*β*42/40 ratio, p-tau 181, p-tau 217, sTREM2, neurofilament light (NfL), glial fibrillary acidic protein (GFAP), apoE, amyloid PET SUVR, MMSE, and FAQ. A large fraction (26 modules, 89.6%) of identified modules showed significant correlations with at least one clinical feature (*P*_FDR_ *<* 0.05). M3 and M18 were significantly associated with neuropathological AD hallmarks, and four modules, including M7, M11 and M16, were linked to cognitive performance. M26 revealed a unique and strong correlation with plasma apoE levels (Fig. 3a, Supplementary Tables 10-11).

To further delineate the correlation between proteomic network modules with diagnostic classification, we compared the module eigenproteins across the AD continuum. A global Kruskal–Wallis rank-sum test (*P*_FDR_ *<* 0.05) identified six modules that varied significantly among four statuses (CN Core1–, CN Core1+, MCI Core1+ and AD dementia Core1+). Post hoc pairwise comparisons revealed that four modules (M1, M11, M20, and M26; *P*_FDR_ *<* 0.05) differed between biologically defined AD and CN Core1–, whereas five modules (M1, M10, M11, M16, and M20; *P*_FDR_ *<* 0.05) distinguished among AD continuum subgroups (Fig. 3b-3g, Supplementary Table 12). We assessed the biological significance of each co-expression module using Gene Ontology (GO) pathway enrichment analysis [21]. Twenty-one of 29 modules were enriched for diverse biological functions, processes, and components (Extended Data Fig. 5, Supplementary Table 13). To investigate the cell-type specificity of modules, we conducted Expression Weighted Celltype Enrichment (EWCE) analysis based on the proportion of mRNA expression of proteins within each module. This analysis revealed significant enrichment of microglial proteins in M16 and M24 and OPC markers in M3 and M6. M7 was uniquely enriched in inhibitory neurons, and astrocyte proteins were enriched in M19 and M27. These findings suggest that GO-defined biological pathways within each module may reflect cell-type-specific alterations relevant to AD (Extended Data Fig. 6, Supplementary Table 14).

### 2.5 Protein importance ranking in AD

To translate our findings into clinically meaningful proteomic panels, we trained Light-GBM classifiers on the DAPs identified for biologically defined AD and for each disease stage, ranking proteins by information gain.

For biologically defined AD versus CN Core1-, 45 proteins together accounted for *>* 85% of cumulative gain, with plasma ACHE ranking highest, followed by LRRN1 and CPLX2 (Fig. 4a). Sequential forward selection showed a sharp rise in AUC after inclusion of the top six proteins (ACHE, LRRN1, CPLX2, C1QTNF5, ALOX5, and LRIG3; AUC = 0.792), which further improved to 0.815 once the next four (IGLL1, GBA, SPC25, and ST8SIA1) were added; performance plateaued thereafter. We therefore retained these ten proteins as the Core1+ proteomic signature for biologically defined AD (Fig. 4a, Supplementary Table 15).

**Fig. 4.**
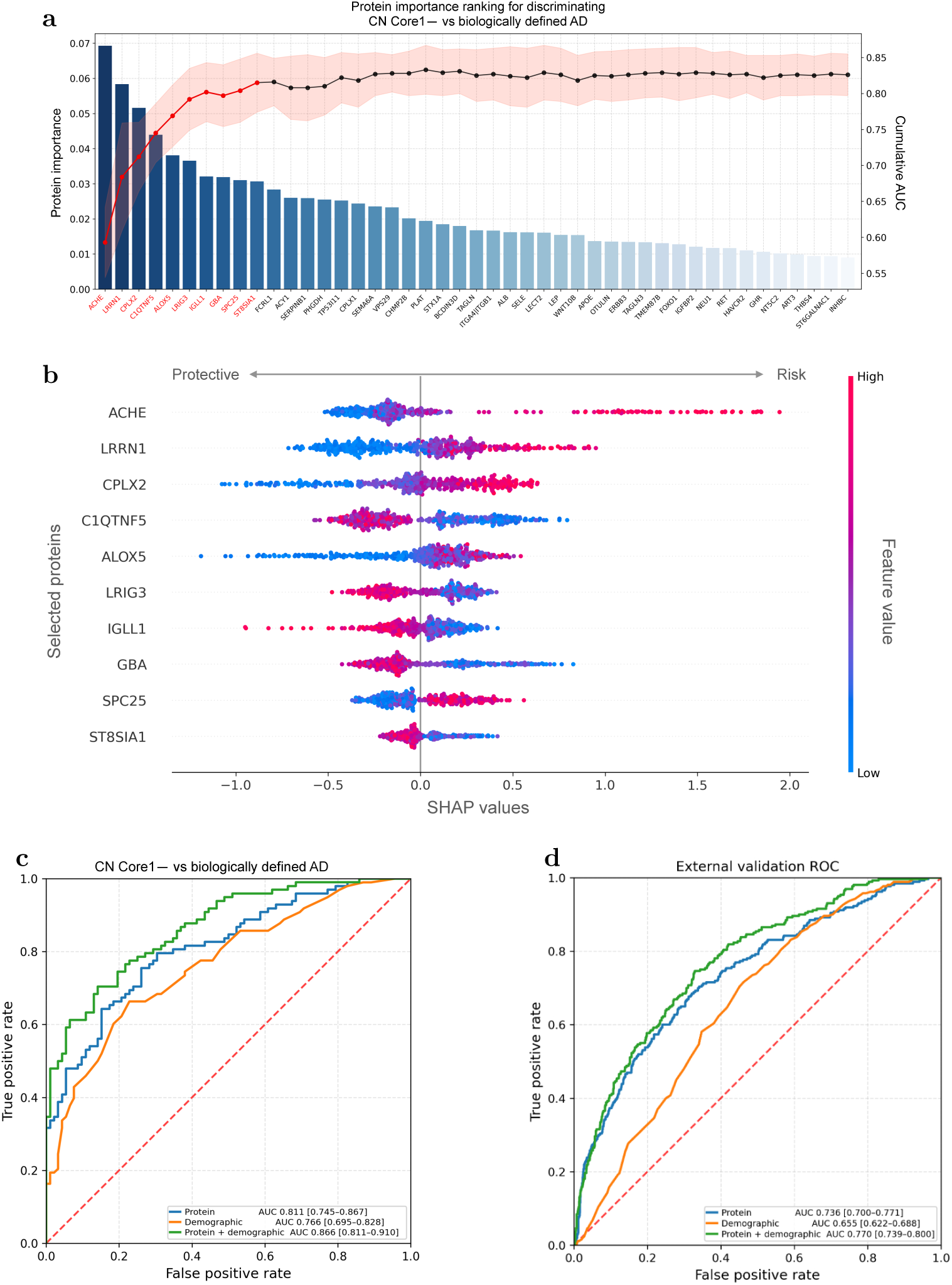
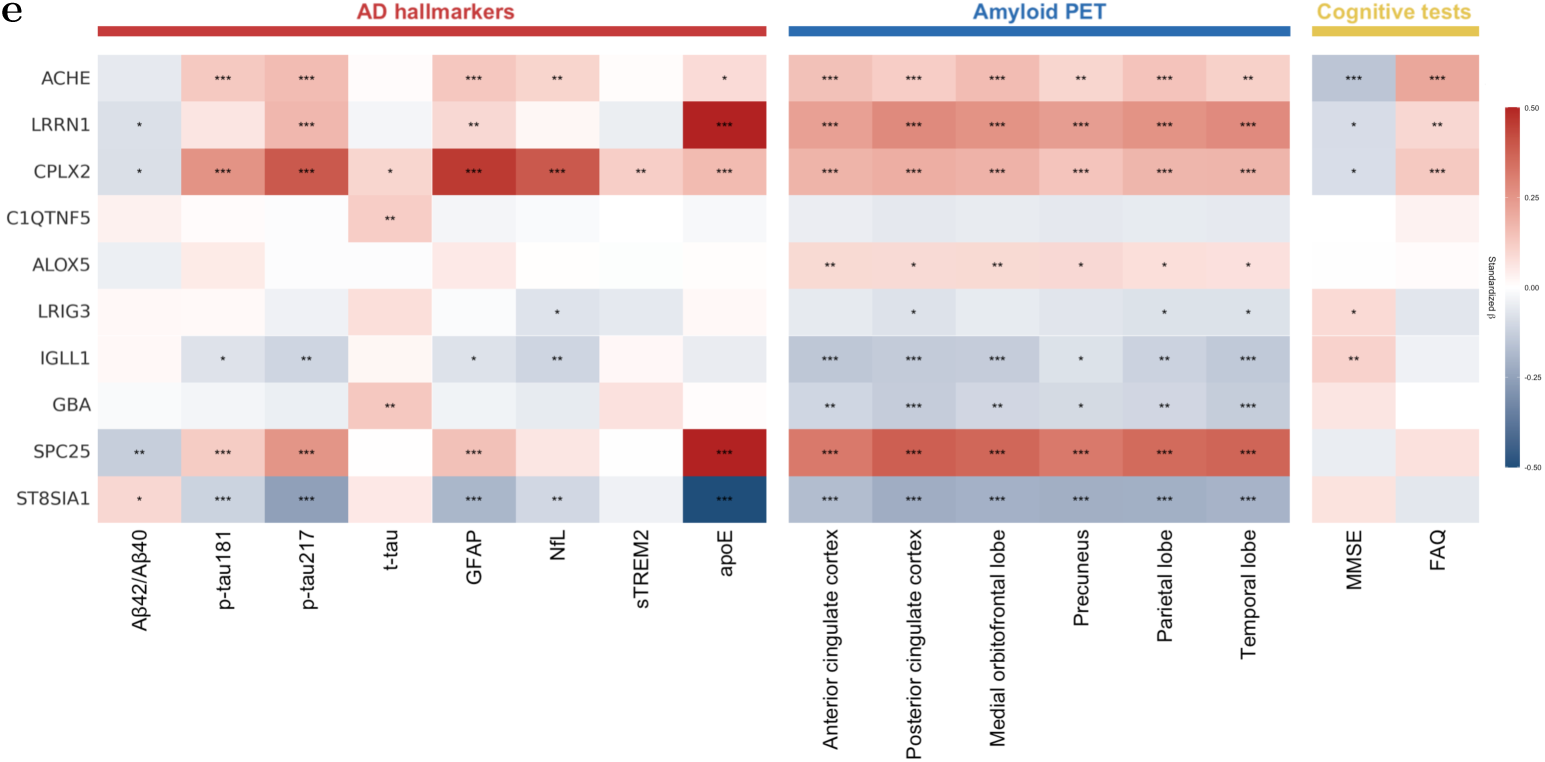
Protein selection, feature importance, and ROC curves for distinguishing biologically defined AD from controls. **a,** Protein panel selection in the derivation set. The bar plot demonstrates ranked importance of proteins according to contribution to distinguishing CN Core1– from biologically defined AD, assessed by information gain. The line plot illustrates the cumulative AUC (right axis) obtained by sequentially adding proteins one at a time in each iteration. Shaded areas represented standard errors estimated using 5-fold cross-validation within the derivation set. Proteins retained in the final panel are highlighted in red. Protein selection was terminated when no further AUC increment *≥*0.005 was observed over two consecutive iterations. **b,** SHAP plot illustrating the contributions of selected proteins to model predictions. The x axis indicates the impact on model output, with positive values (right) corresponding to the higher likelihood of developing AD and negative values (left) corresponding to cognitive normal status. Colour indicates feature value. **c,d,** ROC curves derived in the internal replication cohort (c) and external cohort (GNPC version 1.1 Contributor C) (d) showing discriminatory performance (AUC with 95% CIs) of the selected protein panel alone and combined with demographic variables (age and sex) for distinguishing CN Core1– from biologically defined AD. AUCs and CIs were estimated using bootstrap resampling (2,000 iterations). **e,** Heat map showing associations of the selected proteins with AD endophenotypes. Colours represent beta estimates and asterisks indicate statistical significance after Bonferroni correction (^no symbol^*P ≥* 0.05, * *P <* 0.05, ** *P <* 0.01, *** *P <* 0.001).

SHAP analysis confirmed ACHE as the dominant contributor, spanning the widest range along the model-output axis, while GBA exhibited apparent non-linear effects (Fig. 4b, Supplementary Table 15). To obtain a consistent risk score, we then fitted logistic regression models on the derivation set using the selected panel, demographics (age and sex), and their combination, and evaluated them on the held-out replication set. The ten-protein panel achieved an AUC of 0.811 (95% CI 0.745–0.867); demographics alone yielded 0.766 (95% CI 0.695–0.828); and combining the two improved performance to 0.866 (95% CI 0.811–0.910) (Fig. 4c, Supplementary Tables 16-17).

The panel was further validated in an independent external cohort (GNPC version 1.1 Contributor C) [19, 20]. The logistic regression risk model built on the selected proteins distinguished AD from CN with an AUC of 0.736 (95% CI 0.700–0.771), and the addition of demographics raised this to 0.770 (95% CI 0.739–0.800) (Fig. 4d, Supplementary Table 18).

We next built stage-specific classifiers for MCI Core1+ and AD dementia Core1+, in which 20 and 40 proteins, respectively, accounted for *>* 85% of cumulative gain. In MCI Core1+, CDON ranked highest, followed by CPLX2, AZGP1, and LRRN1 (Extended Data Fig. 7a, Supplementary Table 15); in AD dementia Core1+, ACHE again contributed most, followed by SMOC1, LRRN1, and CPLX2 (Extended Data Fig. 7d, Supplementary Table 15). Sequential forward selection yielded compact panels of nine and twelve proteins, respectively, beyond which AUC plateaued. SHAP plots illustrated the magnitude and direction of each protein’s contribution at both stages (Extended Data Fig. 7b and 7e, Supplementary Table 15). Combining proteins with demographics in logistic regression risk models produced the strongest performance, with AUC = 0.850 (95% CI 0.765–0.923) in MCI Core1+ and AUC = 0.856 (95% CI 0.783–0.919) in AD dementia Core1+, both surpassing demographic-only models (Extended Data Fig. 7c and 7f, Supplementary Tables 16-17).

SHAP values revealed how selected proteins influenced the model discrimination at each disease stage (Extended Data Fig. 7b and 7e, Supplementary Table 15). We then evaluated whether combining plasma proteins with demographics improved stage-specific classification. Combined models achieved the best performance, with AUC = 0.850 (95% CI 0.765–0.923) in MCI Core1+ and AUC = 0.856 (95% CI 0.783–0.919) in AD dementia Core1+, both outperforming models based on demographic information alone (Extended Data Fig. 7c and 7f, Supplementary Tables 16-17).

### 2.6 Associations with AD endophenotypes

We subsequently assessed whether proteins in the selected panel were associated with AD endophenotypes. All ten proteins in the panel were significantly associated with PBBD biomarkers, amyloid PET imaging measured across brain regions, and cognitive assessments.

Within the selected panel, ACHE, LRRN1, CPLX2, IGLL1, SPC25, and ST8SIA1 demonstrated strong associations with p-tau 217, highlighting their relevance to brain amyloid positivity as reflected in the peripheral circulation [22, 23]. LRIG3 was strongly correlated with NfL levels, a marker of neurodegeneration. *APOE-*ε*4* -dependent proteins, including LRRN1, SPC25, and ST8SIA1, exhibited robust correlations with circulating apoE levels. For amyloid neuroimaging biomarkers, nine proteins from the constructed panel were markedly associated with amyloid PET SUVRs, a standardised measure of A*β* burden in the brain [24]. These associations were consistently observed across six brain regions. ACHE, LRRN1, CPLX2, LRIG3, and IGLL1 exhibited significant associations with cognitive performance assessed by MMSE and FAQ, which respectively measure global cognition and functional independence. Pronounced correlations were observed for ACHE, LRRN1, and CPLX2 across all three endophenotype categories (Fig. 4e, Supplementary Table 19).

For the three top-ranking proteins, we further explore their performance in predicting cognitive decline using longitudinal follow-up measurements from GNPC version 1.1, comprised of Contributor B, F and R cohorts with plasma profiling using the Somalogic assay. Cognitive trajectories, assessed by MMSE scores, were derived from linear mixed-effects (LME) models. All three proteins were independently validated and showed significant associations with longitudinal MMSE decline (*P <* 0.05; Extended Data Fig. 8a-8c, Supplementary Table 20).

### 2.7 Genetic evidence linking DAPs with AD

We undertook bidirectional two-sample MR to evaluate evidence for potential causal relationships between DAPs and AD risk. Genome-wide association study (GWAS) for the 69 DAPs defined in biologically defined AD relative to CN Core1– was conducted within Bio-Hermes samples. We used the largest publicly available AD GWAS in the population of European ancestry as the disease dataset (111,326 cases and 677,663 controls) [17].

In the forward direction MR analysis (protein abundance as exposure and AD as outcome), we investigated significant *cis*-acting instrumental variants (IVs; *P <* 5*×*10*^−^*^6^) for each DAP using the genetic-instrument selection strategy and identified 12 DAPs had *cis*-acting IVs. Three proteins (ACHE, FCRL1, and LRIG3) were nominally significantly associated with AD risk (*P <* 0.05; Table 2 and Supplementary Table 21), and the ACHE-AD association remained significant after FDR correction (*P*_FDR_ *<* 0.05; Supplementary Table 21). This result was concordant with dynamic changes in the circulating ACHE levels and provided genetic evidence consistent with a potential role of ACHE in AD pathological progression.

**Table 2.** Bidirectional MR associations between proteins and Alzheimer’s disease.

| Protein | Forward MR (proteins → AD) |  |  | Backward MR (AD → proteins) |  |  | Novel |
| --- | --- | --- | --- | --- | --- | --- | --- |
|  | No. SNPs | Slope (95% CI) | p value | No. SNPs | Slope (95% CI) | p value |  |
| ACHE | 1 | 0.394 (0.154–0.634) | 0.0013 | 52 | 0.011 (–0.029– 0.051) | 0.6004 | No |
| FCRL1 | 1 | –0.321 (–0.603– –0.039) | 0.0256 | 52 | 0.007 (–0.028– 0.042) | 0.6946 | Yes |
| LRIG3 | 2 | –0.706 (–1.345– –0.067) | 0.0303 | 52 | 0.003 (–0.015– 0.020) | 0.7739 | Yes |
| INHBC | NA | NA | NA | 52 | –0.037 (–0.068– –0.005) | 0.0216 | Yes |
| AIMP1 | NA | NA | NA | 52 | –0.021 (–0.040– –0.002) | 0.0332 | Yes |

Note: Wald ratio estimates were used for single-SNP analyses; IVW was used for multiple-SNP analyses. AD, Alzheimer’s disease; CI, confidence interval; SNP, single nucleotide polymorphism.

To explore potential for reverse causation, we performed backward MR analysis (AD as exposure and protein abundance as outcome). Nominally significant associations (*P <* 0.05; Table 2) were observed between AD and two proteins (INHBC and AIMP1) using the random-effects inverse-variance weighted (IVW) model. Additional MR methods, including MR-Egger, weighted median, and weighted mode, yielded consistent effect estimates (Supplementary Table 22). These findings did not pass multiple-testing correction. MR-Egger intercepts indicated no directional pleiotropy, and Cochran’s Q test provided no evidence for heterogeneity, supporting AD as a potential driver of altered abundance for these nominally significant proteins.

To investigate whether the causal associations were driven by the *APOE-*ε*4* isoform, we performed MR using GWAS data calculated with additional adjustment for *APOE-*ε*4* carrier status (model 2) in sensitivity analyses. The ACHE-AD association remained significant, and the MR estimate shifted to the right (*β* = 0.395, *P*_FDR_ *<* 0.05; Supplementary Tables 23-24), providing evidence consistent with an *APOE-*ε*4* -independent association between genetically predicted ACHE abundance and AD risk. These findings were concordant with ACHE as an *APOE-*ε*4* -independent protein, with dysregulated ACHE abundance potentially reflecting upstream processes in AD pathology.

## 3 Methods

### 3.1 Participants and stratification in the Bio-Hermes Study

The Bio-Hermes Study was a multicentre, cross-sectional project conducted across 17 research sites between April 2021 and November 2022 by the Global Alzheimer’s Platform Foundation (GAP). It enrolled participants aged 60 to 85 years from community-based populations in the United States [13]. Participant stratification was based on standard clinical screening procedures that determined clinical presentation as cognitively normal, MCI, or clinically diagnosed mild AD dementia. Inclusion criteria are summarised in Supplementary Table 1.

A total of 988 participants (264 clinically diagnosed AD dementia patients, 308 MCI, and 416 cognitively normal controls) were recruited to establish the plasma proteomics cohort, with demographic data and blood samples collected from all participants. Among the proteomics cohort, 935 participants also underwent brain amyloid PET imaging using the FDA-approved tracer florbetapir F-18 (Amyvid; 18F-AV-45) and were included in the main analyses. All PET scans were centrally interpreted by an expert trained in the manufacturer read process, with final determinations made in accordance with manufacturer standards. PET SUVRs were computed by averaging tracer uptake across a cortical composite of AD-relevant regions (anterior and posterior cingulate, medial orbitofrontal, parietal, precuneus, and temporal regions) and normalising to a reference region (cerebellum).

In accordance with the updated Revised Criteria of Alzheimer’s Association Workgroup for diagnosing and staging AD, amyloid PET imaging was designated as the Core1 imaging biomarker, reflecting AD-specific neuropathological changes and supporting early detection. Amyloid PET results were classified as Core1+ for abnormal scans and Core1– for normal scans. Participants were stratified jointly by clinical diagnosis and Core1 biomarker status as follows: CN Core1– (n = 306), CN Core1+ (n = 81), MCI Core1+ (n = 104), MCI Core1– (n = 183), AD dementia Core1+ (n = 141), non-AD dementia Core1– (n = 95). For analytic purposes, CN Core1+, MCI Core1+ and AD dementia Core1+ were combined into the biologically defined AD group (n = 326).

In the Bio-Hermes study, blood samples were generally collected without requiring participants to fast. Samples were drawn, centrifuged, frozen, and shipped in accordance with assay-specific protocols from the following laboratories: Quanterix for Simoa analysis of A*β*40, A*β*42, A*β*42/A*β*40, NfL, GFAP, t-tau and p-tau181 [25]; C_2_N for the measurement of apoE level; Eli Lilly and Company Clinical Diagnostics Laboratory for measurement of p-tau217 [26].

### 3.2 SomaScan proteomics data processing

Proteomic profiling in the Bio-Hermes Study was performed using the SomaScan v4.1 assay (SomaLogic Inc.). This assay measures 7,584 aptamers targeting approximately 6,400 unique human proteins. The platform uses an aptamer-based approach with slow off-rate modified aptamers (SOMAmers), which incorporate chemically modified nucleotides to bind target proteins with high specificity and affinity. Plasma aptamer abundance was reported in relative fluorescent units (RFU) [27]. Proteomic data were processed using SomaLogic’s standardised adaptive normalisation by maximum likelihood (ANML) pipeline to minimise inter- and intra-plate technical variability [28]. Samples with signal intensities that substantially deviated from expected ranges were flagged by SomaLogic as quality concerns and excluded. After initial quality control, 964 participants remained eligible for inclusion.

Further quality control followed previously reported procedures [11]. RFU pro-teomic values were log10-transformed to approximate a normal distribution. Quality control proceeded in five steps: (1) IQR-based outlier detection using log10-transformed values, with extreme outliers defined relative to the 1st and 3rd quantiles by greater than 2.25 fold the IQR; (2) removal of analytes and samples with *<* 65% call rate; (3) recalculation of analyte call rate and removal of analytes with *<* 85% call rate; (4) recalculation of sample missingness and removal of samples with *<* 85% call rate; (5) exclusion of non-human analytes and analytes without protein targets to ensure biological relevance. After all exclusions, the final Bio-Hermes proteomics ana-lytic matrix comprised 7,289 aptamers and 961 participants, including 405 cognitively normal controls, 305 participants with MCI, and 251 patients with AD.

To screen for potential batch effects, we performed principal component analysis (PCA) using the ‘prcomp’ function in the ‘stats’ R package. Scatter plots of the first two proteomic PCs revealed two distinct batches. After adjustment for proteomic PC1 and PC2, the batch effect was effectively removed (Extended Data Fig. 1). Finally, proteomic aptamer levels were z-score normalised after log10 transformation using the ‘scale’ function in R with both the scale and centre options set to TRUE.

### 3.3 Participants in the GNPC

GNPC version 1.1 consists of 23 independent sites contributing 31,111 plasma proteomic samples from 21,979 individuals with a range of clinical backgrounds, including AD, Parkinson’s disease (PD), frontotemporal dementia (FTD), amyotrophic lateral sclerosis (ALS) and CN controls. Proteomic analysis was conducted using the SomaScan v4.1 assay (SomaLogic Inc.) [11, 19].

GNPC version 1.1 includes three independent longitudinal cohorts (Contributors B, F and R), each of which included individuals who were cognitively normal at baseline and had longitudinal MMSE assessments as well as plasma proteomic data. After applying the same standardised quality control procedures to exclude outlier samples and analytes, the final GNPC proteomics-derived dataset retained 571 donors and 1655 samples and 7,289 aptamers for validation analyses.

GNPC version 1.1 contributor C also included participants assessed by cerebrospinal fluid (CSF) amyloid ratio and amyloid PET status [20, 29]. After applying the same quality control procedures, the final proteomic analytical dataset comprised 7,289 aptamers, and 1,816 donors were included in validation analyses. Participant characteristics are summarised in Supplementary Table 3.

### 3.4 Differential abundance analysis

To identify differentially abundant proteins across AD continuum, we fitted separate multiple linear regression models for each protein. For distinct disease statuses (CN Core1+, MCI Core1+, and AD dementia Core1+), participants were contrasted with CN Core1– as the reference group. Protein abundance (log10-transformed, z-scored RFU) was modelled as the dependent variable, and disease status was modelled as the independent variable, adjusting for age, sex, and the first two proteomic PCs (model 1; four-covariate model). PC1 and PC2 were included as covariates to correct for the batch effects and other potential confounders in the proteomic data (Extended Data Fig. 1). In sensitivity analyses, *APOE* -*ε*4 carrier status (*≥*1 *APOE* -*ε*4 allele) was included as an additional covariate (model 2; five-covariate model). Statistical significance was defined as a Benjamini–Hochberg FDR-adjusted *P* value *<* 0.05.

### 3.5 Cell-type enrichment analysis

To evaluate cell-type expression of genes encoding DAPs, we used single-nucleus transcriptomics data from middle temporal gyrus (MTG) samples from seven postmortem human brains available from the Allen Brain Atlas consortium (https://data.nemoarchive.org/publication_release/Great_Ape_MTG_Analysis/). We downloaded the single-nucleus RNA-sequencing (snRNA-seq) datasets comprising 156,285 high-quality nuclei. The gene expression matrix was normalised using the SCTransform function in the R package Seurat (v.4.4.0) and annotated with cell subclasses available from the Allen Brain data. Subclasses were grouped into six major cell classes: four non-neuronal cell types (astrocytes, OPCs, oligodendrocytes, and microglia) and two neuronal cell types (excitatory glutamatergic neurons and inhibitory GABAergic neurons) [18]. We calculated the average expression from six cell subclasses by applying the AverageExpression function in Seurat and subsequently computed percentage expression across cell types.

We also performed cell-type enrichment analysis using the EWCE R package (v.1.6.0), employing the Allen Brain MTG dataset as the reference [18]. Enrichment was estimated using 10,000 bootstrap resamples, with key gene hits as the target list and the SomaScan-measured proteins as the background. Tested cell types were astrocytes, oligodendrocytes, OPCs, microglia, excitatory glutamatergic neurons and inhibitory GABAergic neurons. Benjamini–Hochberg FDR correction was used to control for multiple testing.

### 3.6 Co-expression module analysis

We constructed a proteomic co-expression network using the R package WGCNA (v.1.73), a robust computational approach for reducing the dimensionality of high-throughput proteomic data and detecting protein modules based on pairwise correlations [30]. Before network construction, protein abundances were adjusted by regressing out age, sex, and the first two proteomic PCs. The resulting standardised residuals were used for subsequent analyses. To focus on biologically defined AD, we included all Core1+ samples (CN Core1+, MCI Core1+, and AD dementia Core1+) along with CN Core1-samples as a control group, while excluding MCI Core1- and non-AD dementia Core1-groups. Using this filtered matrix as input, the optimal soft-thresholding power was set to *β* = 6, determined by balancing scale-free topology fit and mean connectivity. To enhance robustness against outliers, a protein-protein correlation matrix was computed using biweight mid-correlations and served as input for clustering. A signed network was implemented to preserve positive and negative correlations. We calculated the topological overlap matrix (TOM) to quantify inter-connection within network modules and then performed hierarchical clustering based on dissimilarity (1-TOM) to classify proteins into distinct modules [31]. This procedure defined 29 protein modules.

Module eigenproteins were defined as the first principal component of standardised protein abundances within each module, explaining covariance across module proteins. For each module, eigenprotein-based intramodular connectivity *(k* ME) quantified correlation strength between each protein and its module eigenprotein. To avoid aberrant module assignments, a post hoc quality control procedure was applied to flag proteins with weak module membership (*k* ME *<* 0.25). After this step, all proteins remained in their originally defined modules [32, 33]. We computed Pearson correlations between module eigenproteins and AD endophenotypes, and used Kruskal-Wallis rank-sum tests to assess differences in module eigenproteins across AD continuum stages. For both analyses, *P* values were adjusted using the Benjamini–Hochberg FDR method. Network module preservation across cohorts were assessed using *Z*_summary_ composite preservation scores. For each module, *Z*_summary_ composite preservation scores were generated using the network derived from the Bio-Hermes Study proteomics as the reference template and the external cohort as the target dataset, with 500 permutations [32, 33]. We further validated the proteomic architecture using t-distributed stochastic neighbour embedding (t-SNE) [34]. Dimensionality reduction was implemented with R package Rtsne **[35]** and was applied to proteins ranked in the top 25% by *k* ME value within each network module [32, 33]. Proteins were visualized based on their WGCNA-derived module membership, with colors indicating the corresponding module assignments.

### 3.7 Pathway enrichment analysis

Pathway enrichment analysis was conducted using the clusterProfiler R package (v.4.18.2) [36] and the GO database, including biological process (BP), cellular component (CC), and molecular function (MF) annotations[21]. The background set was defined as the SomaScan-measured proteins used across analyses. Statistical significance was determined using the Benjamini–Hochberg correction, and GO terms with *P*_FDR_ *<* 0.05. were retained. To reduce redundancy in displayed results, the ’simplify’ function was applied with a similarity threshold of 0.7.

### 3.8 Machine learning prediction models

AD risk prediction models were implemented in Python (v3.11.13) using Light-GBM (v4.6.0), with hyperparameter optimization performed in Optuna (v4.6.0) and downstream ROC modeling performed in scikit-learn (v1.7.2). The models aimed to distinguish biologically defined AD (Core1+) and stage-specific groups (MCI Core1+ and AD dementia Core1+) from CN Core1–. The cohort was split once into a derivation set (70%) and a held-out replication set (30%).

The pipeline comprised two steps: protein selection and risk-model evaluation. For protein selection, dysregulated proteins from the differential analyses (DAPs) were used as candidate features. All tuning and feature selection were performed exclusively within the derivation set using 5-fold cross-validation, and the replication set was used only for final evaluation. For each contrast (biologically defined AD vs CN Core1–, AD dementia Core1+ vs CN Core1–, and MCI Core1+ vs CN Core1–), Light-GBM hyperparameters were optimized with Optuna and a final LightGBM model was trained on the derivation set. Proteins were ranked using LightGBM gain-based feature importance. Proteins explaining *>* 85% of cumulative gain were retained, and feature importance was re-estimated on this reduced feature set using 5-fold cross-validation on the same derivation set. Sequential forward selection was then applied by adding one protein at a time in the updated ranking and re-fitting models across derivation folds, with cumulative discrimination tracked by AUC. The final panel size was selected at the AUC plateau, defined as two consecutive steps with incremental AUC gain *<* 0.005.

To quantify discriminative performance in a consistent risk-score framework, ROC analyses were conducted using logistic regression, which provides a consistent linear risk score for demographic-only and combined models. Logistic regression models were fitted using the same derivation set, and AUC was evaluated on the replication set and an external cohort (GNPC version 1.1 Contributor C). ROC analyses were performed for the selected protein panel alone and for the panel combined with demographic covariates (age and sex). Differences between AUCs were assessed using DeLong’s test.

### 3.9 Regression models

Multiple linear regression models examined associations between each protein in the constructed panel with AD endophenotypes, including PBBD biomarkers, amyloid PET SUVRs, and cognitive assessments. In each model, protein level was the independent variable predicting specific AD-related outcome. Outcomes from brain-derived biomarkers and PET imaging were natural log-transformed and standardised (*z* -transformation) to facilitate comparability across modalities. Cognitive measures were standardised (*Z* -transformation) only, as log transformation was not appropriate for these variables. All regressions were adjusted for age, sex, and the first two proteomic PCs. Benjamini–Hochberg correction was applied, with statistical significance defined as adjusted *P <* 0.05.

We further evaluated associations between the three top-ranked proteins and longitudinal cognitive performance for CN donors in the validation cohorts (GNPC version 1.1 Contributors B, F and R) using LME models. The outcome was repeatedly measured MMSE score. For each protein, we fitted a model including the interaction between protein level and time (coded as years since baseline, Δage) to capture protein-specific differences in the rate of cognitive change. Models were adjusted for age at baseline, sex, the first two proteomic PCs, and baseline MMSE score.

### 3.10 Genomic data quality control and genome-wide association analysis

Genetic data were obtained for 952 individuals, provided in VCF format and aligned to the hg38 reference genome. Quality control was conducted using PLINK 1.9. Single nucleotide polymorphisms (SNP) with low imputation quality (INFO *≤* 0.7), non-biallelic variants and duplicate SNP names were filtered. Samples were excluded for relatedness (identity by descent, PI HAT *>* 0.2) or high missingness (sample call rate *<* 95%), resulting in 945 individuals. To assess population stratification, the dataset was merged with the 1000 Genomes Project reference panel[37], and PCA was conducted to cluster individuals into three ancestry groups: European and near-European (EUR = 770), African (AFR = 103), and East Asian (ASN = 9). To maximise statistical homogeneity, only individuals identified as European and near-European were retained. The first two genomic PCs are shown in Supplementary Figure 1. SNPs with minor allele frequency (MAF) *<* 5% or deviations from Hardy-Weinberg equilibrium (*P <* 1 *×* 10*^−^*^6^) were removed. After restricting to participants who passed both genotype and proteomics quality control, the final sample comprised 744 individuals. The final set of autosomal genetic variants comprised 5,687,765 SNPs.

SNP-based association analyses were conducted for differentially abundant proteins identified between biologically defined AD and CN Core1– group using linear regression models implemented in PLINK 1.9. Each protein was tested using two models. Model 1 was adjusted for the first five genomic principal components (PC 1-5) and the four covariates used in the differential proteomic analyses (age, sex, proteomics PC 1-2). Model 2 included the same covariates as Model 1, with additional adjustment for *APOE-*ε*4* carrier status used as the sensitivity analysis.

### 3.11 Mendelian randomization

To investigate potential causal associations between the identified DAPs and AD, we performed Bidirectional two-sample MR. For forward MR analysis, we prioritised cis-acting SNPs for each protein. The *cis-*region was defined as a genomic window within 1 Mb upstream or downstream of the transcription start site of the gene encoding the protein of interest. For reverse MR analysis, we used summary statistics from the largest publicly available GWAS of AD in the European ancestry population (111,326 cases and 677,663 controls)[17]. Variants in the major histocompatibility complex region (chromosome 6: 27510020-34480577) and the APOE region (chromosome 19: 43905796-45909393) were excluded due to complex linkage disequilibrium (LD) and high pleiotropy with AD reported previously [38, 39].

To ensure the independence of IVs, LD clumping was performed using reference panel from the 1000 Genomes Project European ancestry population [37], with a 1 Mb clumping window and LD threshold of *r* ² *<* 0.01. Variant selection was restricted to variants with a minor allele frequency (MAF) *>* 5%. For AD GWAS, a significant threshold of *P <* 5 *×* 10*^−^*^8^ was used. For each protein, statistically significant *cis* variants reaching *P <* 5 *×* 10*^−^*^6^ were retrieved. This relaxed threshold was used to maximise inclusion of biologically relevant instruments within the restricted *cis*-regions, and 12 of 69 proteins yielded *cis*-acting IVs with *P <* 5 *×* 10*^−^*^6^. To reduce the risk for weak instrument bias, we calculated the *F* -statistic for each SNP, and we excluded genetic instruments with an *F* -statistic of ¡10 [**?**].

Bidirectional two-sample MR was implemented using the ‘TwoSampleMR’ and ‘MR’ R packages. For forward MR analyses, Wald ratio or random-effects inverse-variance weighted (IVW) methods were used depending on the number of IVs available for each protein. For reverse MR analyses with 52 genetic instruments, random-effects IVW was used as the primary analysis for AD-protein associations due to its high sensitivity for causal inference. Three additional MR methods, including MR-Egger regression, weighted median, and weighted mode, were applied as complementary analyses to enhance the reliability of the results. Directional horizontal pleiotropy was tested using the Egger intercept, and heterogeneous effects among causal estimates were assessed using Cochran’s Q test. Sensitivity analyses used protein GWAS summary statistics derived from models adjusted for *APOE-*ε*4* carrier status (model 2). Benjamini–Hochberg correction was applied, defining statistical significance as *P*_FDR_ *<* 0.05.

## 4 Discussion

We integrated high-throughput plasma proteomic profiling using SomaScan and amyloid PET imaging to quantify AD pathology in the Bio-Hermes Study. Amyloid PET was used as a Core1 biomarker to define the initial stage of the disease, consistent with the 2024 Revised Criteria for Diagnosis and Staging of AD [1]. Our analyses identified differentially abundant plasma proteins across the AD continuum, mapped these proteins to co-expression molecules capturing diverse aspects of AD pathophysiology, and used supervised machine learning to derive protein panels with discriminatory performance for AD status. Integration of genomic data provided additional evidence linking selected proteomic signatures to AD progression.

We identified and characterised multiple plasma proteins that were differentially abundant in clinically diagnosed AD dementia and biologically defined AD. Biologically defined AD, based on amyloid PET biomarkers, captures the underlying pathological features of AD and aligns with intermediate/high ADNPC validated against autopsy [6, 7]. The biologically based definition is consistent with the revised NIA-AA diagnostic criteria [1]. Only 11 proteins overlapped between the clinically diagnosed and biologically defined comparisons, representing a small proportion of both DAP sets. The smaller number of DAPs in biologically defined AD may reflect reduced heterogeneity compared with clinically diagnosed AD dementia, which can include non-AD pathophysiological disorders. By comparing biologically defined AD with CN Core1– controls, identified proteins are more likely to be associated with AD-specific pathogenic mechanisms rather than with non-AD neurodegenerative processes. Top-ranked proteins in biologically defined AD included SPC25, LRRN1, ACHE, CPLX2, S100A13, BCDIN3D, ST8SIA1, and TBCA. These data provide preliminary evidence that neuropathology-based classification can improve the biological specificity of plasma proteomic signatures.

This study maps blood-based proteomic profiles across the spectrum from CN core1+ and MCI Core1+ to AD dementia Core1+. We identified eight overlapping proteins whose plasma abundances were dysregulated across the AD continuum, from preclinical and prodromal phases to AD dementia. These proteins included established AD-related biomarkers such as NEFL and targets reported in recent proteomics studies, including SPC25, LRRN1, TBCA, and S100A13 [11, 15, 16, 40, 41]. Candidate proteins included BCDIN3D, involved in RNA methylation [42]; ST8SIA1, involved in ganglioside biosynthesis [43]; and FOXO1, involved in insulin-resistance signalling [44]. Because this protein subset was already altered in preclinical AD, these proteins warrant further evaluation as candidate markers of early pathological change. We also identified proteins associated with transition to cognitive impairment (MCI Core1+ and AD dementia Core1+), including the pseudophosphatase PTPRU [45], the neddylation factor DCUN1D5 [46], and the synaptic-vesicle exocytosis regulator CPLX2 [47]. Together with results from recent independent studies [46, 48], these findings can be used to support molecular staging of AD, and prioritise candidates for mechanistic validation and novel therapeutic targets.

Stage-specific protein signatures shifted as disease progressed, encompassing proteins dysregulated transiently during the preclinical phase, proteins emerging with MCI, and proteins altered predominantly in AD dementia. A notable finding was elevation of plasma ACHE that was specific to the AD dementia Core1+ phase. The result remained robust when the analysis was restricted to participants naive to cholinesterase inhibitors, suggesting that ACHE enrichment may precede pharmacological intervention and reflect intrinsic disease-related processes. ACHE is a primary enzyme, that forms a part of the parasympathetic nervous system and is significantly involved in synaptic plasticity and neurotransmission efficiency [49, 50]. It has been repeatedly identified as a key protein in the CSF and brain tissue of patients with AD dementia [51], and cholinesterase inhibitors have long been used to alleviate cognitive and functional symptoms prior to the recent emergence of disease-modifying treatments [52, 53]. Additional evidence suggests broader pathogenic roles for ACHE in AD, including effects on calcium dysregulation, excitotoxicity, and demyelination [49, 54, 55]. Our analyses suggest that molecular transitions across the AD continuum may correspond to distinct stages of cognitive decline, with many DAPs showing stage specificity.

By comparing statistical models with and without *APOE-*ε*4* carrier status adjustment, we categorised proteins as *APOE-*ε*4* -independent or *APOE-*ε*4* -dependent. In AD dementia Core1+, we identified 20 *APOE-*ε*4* -independent and 91 *APOE-*ε*4* -dependent DAPs. In MCI Core1+, 83 detected DAPs were attenuated after *APOE-*ε*4* adjustment. This stratification provides a compartmentalised view of AD-related proteomic signatures and underscores the extent to which the proteomic landscape is shaped by genetic susceptibility and disease stage, a critical consideration for biomarker discovery. *APOE-*ε*4* -independent DAPs, such as ACHE, CPLX2 and SPON1, were predominantly dysregulated in late-stage pathology (MCI Core1+ or AD dementia Core1+). By contrast, the eight overlapping proteins altered across the AD continuum were consistently *APOE-*ε*4* -dependent. Previous studies linked incident AD to plasma levels of *APOE-*ε*4* –dependent proteins including NEFL, LRRN1, TBCA and S100A13 [15], and reported associations between *APOE* genotype and biofluid (plasma and CSF) levels of LRRN1, SPC25, TBCA and FOXO1, although direct associations with AD were not described in this study [16]. Collectively, *APOE-*ε*4* - dependent DAPs may reflect the pathogenic mechanisms driven by *APOE-*ε*4* genotype and provide candidate plasma-based indicators relevant to AD patients carrying this risk allele [56, 57]. We further detected that sex-related protein differences in biologically defined AD. 26 proteins were FDR significant in females compared to four proteins in males. This difference may reflect the limited sample size in subgroup analyses.

By leveraging plasma proteomics alongside multilevel data, including cognitive assessments, blood-based biomarkers, and PET imaging, a systems-level approach was applied to investigate the multifaceted pathophysiology and heterogeneous biological networks underlying the AD continuum. Correlation network and pathway enrichment analyses identified 29 protein co-expression modules [21, 30], 26 of which were linked to AD endophenotypes. Six modules showed stage-specific patterns, demonstrating how pathway signatures map to AD progression. Networks correlated with AD pathogenesis were enriched for biological processes and functions related to synaptic signalling, endosomes, histone binding, extracellular matrix, sulfur binding, chaperone, cytoskeleton, RNA binding/splicing, chemokine activities, nucleotide binding and metabolism, axon development, and cadherin binding pathways.

Decreased abundance of M26 was observed across the AD continuum, suggesting that proteins within this module may help characterise disease progression. M26 was enriched for *APOE-*ε*4* -dependent DAPs, including NEFL, BCDIN3D, ST8SIA1, and TP53I11, and showed a robust correlation with circulating apoE levels, indicating a potential *APOE-*ε*4* -driven contribution to disease pathogenesis. M6 correlated strongly with GFAP, an established marker of inflammation in AD, and was enriched for axon-development pathways and OPC signatures, suggesting that M6 may reflect OPC-linked biological processes related to AD neuroinflammation. M6 contained ACHE, the top *APOE-*ε*4* -independent DAP, as a hub protein. Evidence that dysregulated ACHE impairs OPC differentiation and contributes to demyelination supports a potential role for ACHE in neuroinflammatory mechanisms relevant to AD [58, 59]. M3 correlated strongly with late-stage PBBD biomarkers, including p-tau181, p-tau217, GFAP, and sTREM2. This module contained CPLX2 and several proteins observed exclusively in AD dementia Core1+ (SMOC1, TAGLN, and SPON1), and was enriched for signalling receptor regulator activity and OPC markers. Research showing OPC senescence in the Aβ-plaque environment may provide insight into the roles of OPC-enriched modules in AD [60]. Findings for SMOC1 and SPON1 were consistent with previous proteomic studies [9, 61–63]. Remarkably, SMOC1 expressed within the OPC-related module aligned with recent observations in the BioFinder cohort [61]. In contrast, CPLX2 was less commonly reported and is nominated here as a candidate biomarker for further validation.

Dysregulated proteins were used to develop classification models for biologically defined AD and distinct stages across the AD continuum. We applied LightGBM classifiers and derived plasma protein panels with discriminatory performance. Integrated models combining selected proteins with demographic variables were prioritised to optimise risk prediction. The ten-protein panel for biologically defined AD achieved AUC = 0.815 and outperformed demographic variables in distinguishing all Core1+ groups from CN Core1*−*. Performance significantly improved when the protein panel was combined with demographic variables, and findings were evaluated in an independent cohort. These results suggest that selected proteins are informative candidate markers for non-invasive screening approaches, but prospective clinical validation is required before clinical utility can be inferred. Consistent with WGCNA analyses, selected proteins were distributed across multiple co-expression modules, suggesting that strong predictive performance may reflect complementary information from diverse AD-related biological pathways. ACHE, LRRN1, and CPLX2 were the top three selected proteins, with ACHE contributing most to predictive value and showing associations with AD clinical features.

Stage-specific panels for MCI Core1+ and AD dementia Core1+ demonstrated high discriminatory performance. For MCI Core1+, top predictive proteins were CDON, CPLX2, AZGP1, and LRRN1. For AD dementia Core1+, ACHE had the highest importance gain, followed by SMOC1, LRRN1, and CPLX2. The two stage-specific panels shared selected proteins, suggesting overlapping signatures across the AD trajectory. Shifts in importance ranking between the LightGBM models further illustrate heterogeneity across AD stages. Compared with existing plasma-based biomarkers, this approach has two potential advantages: (1) selected proteins represent pathways implicated in AD-associated dysfunction, as supported by WGCNA analyses; (2) as emerging therapies target A*β* pathology, complementary predictive models not solely reliant on amyloid-related markers may be informative.

MR analyses highlighted genetic evidence consistent with a potential effect of ACHE on AD. This association was directionally consistent across MR methods and supported by differential abundance analysis. ACHE ranked highly for distinguishing biologically defined AD and AD dementia Core1+ exhibited strong associations with cognitive decline in an external cohort and remained significant after FDR correction in MR analyses. These convergent findings support ACHE as a biologically informative candidate biomarker, and while functional and prospective studies are required to establish clinical and mechanistic relevance the result suggests a disease relationship outside of the drug treatment [52, 53]. MR also suggested that genetically predicted levels of FCRL1 and LRIG3 were nominally associated with AD risk, although these associations did not remain significant after FDR correction. ACHE has previously been implicated in AD risk by MR studies[64], whereas FCRL1 and LRIG3 emerged as candidate biomarkers in the Bio-Hermes Study. Bidirectional MR suggested that AD liability may influence abundance of INHBC and AIMP1. Although these associations did not survive FDR correction, the links between AD susceptibility and INHBC or AIMP1 warrant further investigation.

Our study has several limitations. First, although amyloid PET was assessed, tau PET was not measured contemporaneously with baseline plasma proteomics in Bio-Hermes. We therefore could not define biological stages based on tau pathology or investigate proteomic signatures across different levels of tau burden. Incorporating tau PET in future studies will enable finer mapping between proteomic changes and disease progression and will help evaluate whether identified proteins discriminate disease stages longitudinally. Second, although a broad plasma protein panel was quantified, the assay platform does not fully capture the human proteome or comprehensively assess post-translational modifications. The platform’s design may also introduce measurement bias toward secreted proteins, limiting representation of other biologically relevant protein classes. Finally, findings were not validated across multiple proteomic platforms due to technical constraints, and the assay measures relative protein abundance through aptamer binding rather than absolute protein concentrations. However, consistency with previous studies and across independent cohorts support the robustness of the main analyses.

Taken together, this study delineates biologically anchored plasma proteomic alterations across the AD continuum in the Bio-Hermes cohort, identifies co-expression modules linked to AD endophenotypes, and nominates candidate biomarker panels for further validation. Among individual candidates, ACHE, CPLX2, and LRRN1 emerged as informative proteins with associations across model-based classification, endophenotypes and longitudinal cognition.

## Data availability

The Bio-Hermes dataset can be accessed through the Alzheimer’s Disease Data Initiative portal at https://discover.alzheimersdata.org/catalogue/datasets/8395cc96-f249-40db-beaa-4668feb3cc5e. The GNPC version 1.1 harmonised data set (HDS) request link can be found on the GNPC website (https://www.neuroproteome.org/harmonized-data-set-hds).

## Acknowledgments

The authors would like to thank the patients who contributed to the studies used in the manuscript. L.W is funded by Alzheimer’s Research UK (ARUK-RF2020A-005, ARUK-SRF2023B-007). R.M and B.E are funded by Michael J. Fox Foundation (MJFF-022845). Support was provided by Global Alzheimer’s Platform, Race Against Dementia, Gates Ventures, Alzheimer’s Disease Data Initiative.

## Author contributions statement

H.Z. and L.W. conceived the study. H.Z. was responsible for methodology and formal analysis. T.Z., R.M., G.L., and D.L. analysed data. H.Z, B.E., and L.W. prepared the original draft. K.M., T.Q., I.K., V.E.P., S.J. and A.N.H. advised on interpretation of data. All authors reviewed the manuscript.

## Competing interests

A.N.H. receives research funding from GSK and acts as an expert consultant to Scripta Therapeutics. S.J. is shareholder of Oxford Vacmedix Ltd.

## Additional information

Data described in this study are available on application to the Alzheimer’s Disease Data Initiative portal.

## References

[1] Jack Jr, C.R., Andrews, J.S., Beach, T.G., Buracchio, T., Dunn, B., Graf, A., Hansson, O., Ho, C., Jagust, W., McDade, E., et al.: Revised criteria for diagnosis and staging of alzheimer’s disease: Alzheimer’s association workgroup. Alzheimer’s & Dementia 20(8), 5143–5169 (2024)

[2] Zhang, Y., Chen, H., Li, R., Sterling, K., Song, W.: Amyloid *β*-based therapy for alzheimer’s disease: challenges, successes and future. Signal transduction and targeted therapy 8(1), 248 (2023)

[3] Ferrari, C., Sorbi, S.: The complexity of alzheimer’s disease: an evolving puzzle. Physiological reviews 101(3), 1047–1081 (2021)

[4] Jack Jr, C.R., Bennett, D.A., Blennow, K., Carrillo, M.C., Dunn, B., Haeberlein, S.B., Holtzman, D.M., Jagust, W., Jessen, F., Karlawish, J., et al.: Nia-aa research framework: toward a biological definition of alzheimer’s disease. Alzheimer’s & dementia 14(4), 535–562 (2018)

[5] Maheux, E., Koval, I., Ortholand, J., Birkenbihl, C., Archetti, D., Bouteloup, V., Epelbaum, S., Dufouil, C., Hofmann-Apitius, M., Durrleman, S.: Forecasting individual progression trajectories in alzheimer’s disease. Nature Communications 14(1), 761 (2023)

[6] La Joie, R., Ayakta, N., Seeley, W.W., Borys, E., Boxer, A.L., DeCarli, C., Doŕe, V., Grinberg, L.T., Huang, E., Hwang, J.-H., et al.: Multisite study of the relationships between antemortem [11c] pib-pet centiloid values and postmortem measures of alzheimer’s disease neuropathology. Alzheimer’s & Dementia 15(2), 205–216 (2019)

[7] Clark, C.M., Pontecorvo, M.J., Beach, T.G., Bedell, B.J., Coleman, R.E., Doraiswamy, P.M., Fleisher, A.S., Reiman, E.M., Sabbagh, M.N., Sadowsky, C.H., et al.: Cerebral pet with florbetapir compared with neuropathology at autopsy for detection of neuritic amyloid-*β* plaques: a prospective cohort study. The Lancet Neurology 11(8), 669–678 (2012)

[8] Villemagne, V.L., Doŕe, V., Burnham, S.C., Masters, C.L., Rowe, C.C.: Imaging tau and amyloid-*β* proteinopathies in alzheimer disease and other conditions. Nature Reviews Neurology 14(4), 225–236 (2018)

[9] Guo, Y., Chen, S.-D., You, J., Huang, S.-Y., Chen, Y.-L., Zhang, Y., Wang, L.-B., He, X.-Y., Deng, Y.-T., Zhang, Y.-R., et al.: Multiplex cerebrospinal fluid proteomics identifies biomarkers for diagnosis and prediction of alzheimer’s disease. Nature human behaviour 8(10), 2047–2066 (2024)

[10] Yarbro, J.M., Shrestha, H.K., Wang, Z., Zhang, X., Zaman, M., Chu, M., Wang, X., Yu, G., Peng, J.: Proteomic landscape of alzheimer’s disease: emerging technologies, advances and insights (2021–2025). Molecular neurodegeneration 20(1), 83 (2025)

[11] Ali, M., Erabadda, B., Chen, Y., Xu, Y., Gong, K., Liu, M., Pichet Binette, A., Timsina, J., Western, D., Yang, C., et al.: Shared and disease-specific pathways in frontotemporal dementia and alzheimer’s and parkinson’s diseases. Nature medicine 31(8), 2567–2577 (2025)

[12] Heo, G., Xu, Y., Wang, E., Ali, M., Oh, H.S.-H., Moran-Losada, P., Anastasi, F., Gonźalez Escalante, A., Puerta, R., Song, S., et al.: Large-scale plasma proteomic profiling unveils diagnostic biomarkers and pathways for alzheimer’s disease. Nature aging 5(6), 1114–1131 (2025)

[13] Mohs, R.C., Beauregard, D., Dwyer, J., Gaudioso, J., Bork, J., MaGee-Rodgers, T., Key, M.N., Kerwin, D.R., Hughes, L., Cordell, C.B., et al.: The bio-hermes study: biomarker database developed to investigate blood-based and digital biomarkers in community-based, diverse populations clinically screened for alzheimer’s disease. Alzheimer’s & Dementia 20(4), 2752–2765 (2024)

[14] Hussain, Z., Ng, D., Leighton, S., Deligianni, F., Ritchie, C., Cavanagh, J.: Plasma inflammatory biomarker profiles across the alzheimer’s disease spectrum in the bio-hermes cohort. Alzheimer’s & Dementia 22(3), 71257 (2026)

[15] Frick, E.A., Emilsson, V., Jonmundsson, T., Steindorsdottir, A.E., Johnson, E.C., Puerta, R., Dammer, E.B., Shantaraman, A., Cano, A., Boada, M., et al.: Serum proteomics reveal apoe-*ε*4-dependent and apoe-*ε*4-independent protein signatures in alzheimer’s disease. Nature Aging 4(10), 1446–1464 (2024)

[16] Shvetcov, A., Johnson, E.C., Winchester, L.M., Walker, K.A., Wilkins, H.M., Thompson, T.G., Rothstein, J.D., Krish, V., Imam, F.B., (GNPC), G.N.P.C., et al.: Apoe *ε*4 carriers share immune-related proteomic changes across neurodegenerative diseases. Nature Medicine 31(8), 2590–2601 (2025)

[17] Bellenguez, C., Küçükali, F., Jansen, I.E., Kleineidam, L., Moreno-Grau, S., Amin, N., Naj, A.C., Campos-Martin, R., Grenier-Boley, B., Andrade, V., et al.: New insights into the genetic etiology of alzheimer’s disease and related dementias. Nature genetics 54(4), 412–436 (2022)

[18] Siletti, K., Hodge, R., Mossi Albiach, A., Lee, K.W., Ding, S.-L., Hu, L., Lönnerberg, P., Bakken, T., Casper, T., Clark, M., et al.: Transcriptomic diversity of cell types across the adult human brain. Science 382(6667), 7046 (2023)

[19] Imam, F., Saloner, R., Vogel, J.W., Krish, V., Abdel-Azim, G., Ali, M., An, L., Anastasi, F., Bennett, D., Pichet Binette, A., et al.: The global neurodegeneration proteomics consortium: biomarker and drug target discovery for common neurodegenerative diseases and aging. Nature medicine 31(8), 2556–2566 (2025)

[20] Palmqvist, S., Janelidze, S., Stomrud, E., Zetterberg, H., Karl, J., Zink, K., Bittner, T., Mattsson, N., Eichenlaub, U., Blennow, K., et al.: Performance of fully automated plasma assays as screening tests for alzheimer disease–related *β*-amyloid status. JAMA neurology 76(9), 1060–1069 (2019)

[21] Ashburner, M., Ball, C.A., Blake, J.A., Botstein, D., Butler, H., Cherry, J.M., Davis, A.P., Dolinski, K., Dwight, S.S., Eppig, J.T., et al.: Gene ontology: tool for the unification of biology. Nature genetics 25(1), 25–29 (2000)

[22] Ashton, N.J., Brum, W.S., Di Molfetta, G., Benedet, A.L., Arslan, B., Jonaitis, E., Langhough, R.E., Cody, K., Wilson, R., Carlsson, C.M., et al.: Diagnostic accuracy of a plasma phosphorylated tau 217 immunoassay for alzheimer disease pathology. JAMA neurology 81(3), 255–263 (2024)

[23] Asken, B.M., DeSimone, J.C., Wang, W.-E., McFarland, K.N., Arias, F., Levy, S.-A., Fiala, J., Velez-Uribe, I., Mayrand, R.P., Sawada, L.O., et al.: Plasma p-tau217 concordance with amyloid pet among ethnically diverse older adults. Alzheimer’s & dementia: diagnosis, assessment & disease monitoring 16(3), 12617 (2024)

[24] Kněsaurek, K., Warnock, G., Kostakoglu, L., Burger, C., et al.: Comparison of standardized uptake value ratio calculations in amyloid positron emission tomography brain imaging. World journal of nuclear medicine 17(1), 21 (2018)

[25] Ding, X., Zhang, S., Jiang, L., Wang, L., Li, T., Lei, P.: Ultrasensitive assays for detection of plasma tau and phosphorylated tau 181 in alzheimer’s disease: a systematic review and meta-analysis. Translational neurodegeneration 10(1), 10 (2021)

[26] Palmqvist, S., Janelidze, S., Quiroz, Y.T., Zetterberg, H., Lopera, F., Stomrud, E., Su, Y., Chen, Y., Serrano, G.E., Leuzy, A., et al.: Discriminative accuracy of plasma phospho-tau217 for alzheimer disease vs other neurodegenerative disorders. Jama 324(8), 772–781 (2020)

[27] Gold, L., Ayers, D., Bertino, J., Bock, C., Bock, A., Brody, E., Carter, J., Cunningham, V., Dalby, A., Eaton, B., et al.: Aptamer-based multiplexed proteomic technology for biomarker discovery. Nature Precedings, 1–1 (2010)

[28] Candia, J., Cheung, F., Kotliarov, Y., Fantoni, G., Sellers, B., Griesman, T., Huang, J., Stuccio, S., Zingone, A., Ryan, B.M., et al.: Assessment of variability in the somascan assay. Scientific reports 7(1), 14248 (2017)

[29] Barthélemy, N.R., Salvadó, G., Schindler, S.E., He, Y., Janelidze, S., Collij, L.E., Saef, B., Henson, R.L., Chen, C.D., Gordon, B.A., et al.: Highly accurate blood test for alzheimer’s disease is similar or superior to clinical cerebrospinal fluid tests. Nature medicine 30(4), 1085–1095 (2024)

[30] Langfelder, P., Horvath, S.: Wgcna: an r package for weighted correlation network analysis. BMC bioinformatics 9(1), 559 (2008)

[31] Shen, Y., Timsina, J., Heo, G., Beric, A., Ali, M., Wang, C., Yang, C., Wang, Y., Western, D., Liu, M., et al.: Csf proteomics identifies early changes in autosomal dominant alzheimer’s disease. Cell 187(22), 6309–6326 (2024)

[32] Johnson, E.C., Dammer, E.B., Duong, D.M., Ping, L., Zhou, M., Yin, L., Hig-ginbotham, L.A., Guajardo, A., White, B., Troncoso, J.C., et al.: Large-scale proteomic analysis of alzheimer’s disease brain and cerebrospinal fluid reveals early changes in energy metabolism associated with microglia and astrocyte activation. Nature medicine 26(5), 769–780 (2020)

[33] Johnson, E.C., Carter, E.K., Dammer, E.B., Duong, D.M., Gerasimov, E.S., Liu, Y., Liu, J., Betarbet, R., Ping, L., Yin, L., et al.: Large-scale deep multi-layer analysis of alzheimer’s disease brain reveals strong proteomic disease-related changes not observed at the rna level. Nature neuroscience 25(2), 213–225 (2022)

[34] Rangaraju, S., Dammer, E.B., Raza, S.A., Rathakrishnan, P., Xiao, H., Gao, T., Duong, D.M., Pennington, M.W., Lah, J.J., Seyfried, N.T., et al.: Identification and therapeutic modulation of a pro-inflammatory subset of disease-associated-microglia in alzheimer’s disease. Molecular neurodegeneration 13(1), 24 (2018)

[35] Van Der Maaten, L.: Accelerating t-sne using tree-based algorithms. The journal of machine learning research 15(1), 3221–3245 (2014)

[36] Yu, G., Wang, L.-G., Han, Y., He, Q.-Y.: clusterprofiler: an r package for comparing biological themes among gene clusters. Omics: a journal of integrative biology 16(5), 284–287 (2012)

[37] Consortium, .G.P., et al.: A map of human genome variation from population scale sequencing. Nature 467(7319), 1061 (2010)

[38] Pietzner, M., Wheeler, E., Carrasco-Zanini, J., Cortes, A., Koprulu, M., Wörheide, M.A., Oerton, E., Cook, J., Stewart, I.D., Kerrison, N.D., et al.: Mapping the proteo-genomic convergence of human diseases. Science 374(6569), 1541 (2021)

[39] Ferkingstad, E., Sulem, P., Atlason, B.A., Sveinbjornsson, G., Magnusson, M.I., Styrmisdottir, E.L., Gunnarsdottir, K., Helgason, A., Oddsson, A., Halldorsson, B.V., et al.: Large-scale integration of the plasma proteome with genetics and disease. Nature genetics 53(12), 1712–1721 (2021)

[40] Zhang, Y., Guo, Y., He, Y., You, J., Zhang, Y., Wang, L., Chen, S., He, X., Yang, L., Huang, Y., et al.: Large-scale proteomic analyses of incident alzheimer’s disease reveal new pathophysiological insights and potential therapeutic targets. Molecular Psychiatry 30(6), 2347–2361 (2025)

[41] Lu, L., Binette, A.P., Hristovska, I., Janelidze, S., Smets, B., Mayoral, I.C., Vasanthakumar, A., Milkovich, B., Ossenkoppele, R., Krish, V., et al.: Proteomic signatures of the apoe *ε*4 and apoe *ε*2 genetic variants and alzheimer’s disease. medRxiv (2025)

[42] Xhemalce, B., Robson, S.C., Kouzarides, T.: Human rna methyltransferase bcdin3d regulates microrna processing. Cell 151(2), 278–288 (2012)

[43] Mabe, N.W., Huang, M., Dalton, G.N., Alexe, G., Schaefer, D.A., Geraghty, A.C., Robichaud, A.L., Conway, A.S., Khalid, D., Mader, M.M., et al.: Transition to a mesenchymal state in neuroblastoma confers resistance to anti-gd2 antibody via reduced expression of st8sia1. Nature cancer 3(8), 976–993 (2022)

[44] Du, S., Zheng, H.: Role of foxo transcription factors in aging and age-related metabolic and neurodegenerative diseases. Cell & bioscience 11(1), 188 (2021)

[45] Hay, I.M., Fearnley, G.W., Rios, P., Köhn, M., Sharpe, H.J., Deane, J.E.: The receptor ptpru is a redox sensitive pseudophosphatase. Nature Communications 11(1), 3219 (2020)

[46] Mei, Z., Liu, J., Bennett, D.A., Seyfried, N., Wingo, A.P., Wingo, T.S.: Unraveling sex differences in alzheimer’s disease and related endophenotypes with brain proteomes. Alzheimer’s & Dementia 21(5), 70206 (2025)

[47] Komatsu, H., Kakehashi, A., Nishiyama, N., Izumi, N., Mizuguchi, S., Yamano, S., Inoue, H., Hanada, S., Chung, K., Wei, M., et al.: Complexin-2 (cplx2) as a potential prognostic biomarker in human lung high grade neuroendocrine tumors. Cancer Biomarkers 13(3), 171–180 (2013)

[48] Xu, X.-w., Zhou, X.-w., Zhang, L., Wang, Q., Wang, X.-x., Jin, Y.-m., Li, L.-l., Jin, M.-f., Wu, H.-y., Ding, X., et al.: Complexin 2 contributes to the protective effect of nad+ on neuronal survival following neonatal hypoxia-ischemia. Acta Pharmacologica Sinica 46(9), 2363–2375 (2025)

[49] Behl, T., Kaur, I., Sehgal, A., Singh, S., Sharma, N., Gupta, S., Albratty, M., Najmi, A., Alhazmi, H.A., Bungau, S.: Ache as a spark in the alzheimer’s blaze–antagonizing effect of a cyclized variant. Ageing research reviews 83, 101787 (2023)

[50] Lombardo, S., Maskos, U.: Role of the nicotinic acetylcholine receptor in alzheimer’s disease pathology and treatment. Neuropharmacology 96, 255–262 (2015)

[51] Garćıa-Aylĺon, M.-S., Small, D.H., Avila, J., Śaez-Valero, J.: Revisiting the role of acetylcholinesterase in alzheimer’s disease: cross-talk with p-tau and *β*-amyloid. Frontiers in molecular neuroscience 4, 22 (2011)

[52] Giacobini, E.: Long-term stabilizing effect of cholinesterase inhibitors in the therapy of alzheimer’disease. Ageing and Dementia Current and Future Concepts, 181–187 (2002)

[53] Kaduszkiewicz, H., Zimmermann, T., Beck-Bornholdt, H.-P., Bussche, H.: Cholinesterase inhibitors for patients with alzheimer’s disease: systematic review of randomised clinical trials. Bmj 331(7512), 321–327 (2005)

[54] Greenfield, S.A., Day, T., Mann, E.O., Bermudez, I.: A novel peptide modulates *α*7 nicotinic receptor responses: Implications for a possible trophic-toxic mechanism within the brain. Journal of neurochemistry 90(2), 325–331 (2004)

[55] Greenfield, S.A., Cole, G.M., Coen, C.W., Frautschy, S., Singh, R.P., Mekkittikul, M., Garcia-Ratés, S., Morrill, P., Hollings, O., Passmore, M., et al.: A novel process driving alzheimer’s disease validated in a mouse model: therapeutic potential. Alzheimer’s & Dementia: Translational Research & Clinical Interventions 8(1), 12274 (2022)

[56] Frisoni, G.B., Manfredi, M., Geroldi, C., Binetti, G., Zanetti, O., Bianchetti, A., Trabucchi, M.: The prevalence of apoe-*ε*4 in alzheimer’s disease is age dependent. Journal of Neurology, Neurosurgery & Psychiatry 65(1), 103–106 (1998)

[57] Gharbi-Meliani, A., Dugravot, A., Sabia, S., Regy, M., Fayosse, A., Schnitzler, A., Kivimäki, M., Singh-Manoux, A., Dumurgier, J.: The association of apoe *ε*4 with cognitive function over the adult life course and incidence of dementia: 20 years follow-up of the whitehall ii study. Alzheimer’s research & therapy 13(1), 5 (2021)

[58] Cui, X., Guo, Y.-e., Fang, J.-h., Shi, C.-j., Suo, N., Zhang, R., Xie, X.: Donepezil, a drug for alzheimer’s disease, promotes oligodendrocyte generation and remyelination. Acta Pharmacologica Sinica 40(11), 1386–1393 (2019)

[59] Ravichandar, R., Gadelkarim, F., Muthaiah, R., Glynos, N., Murlanova, K., Rai, N.K., Saraswat, D., Polanco, J.J., Dutta, R., Pal, D., et al.: Dysregulated cholinergic signaling inhibits oligodendrocyte maturation following demyelination. Journal of Neuroscience 44(28) (2024)

[60] Zhang, P., Kishimoto, Y., Grammatikakis, I., Gottimukkala, K., Cutler, R.G., Zhang, S., Abdelmohsen, K., Bohr, V.A., Misra Sen, J., Gorospe, M., et al.: Senolytic therapy alleviates a*β*-associated oligodendrocyte progenitor cell senescence and cognitive deficits in an alzheimer’s disease model. Nature neuroscience 22(5), 719–728 (2019)

[61] Pichet Binette, A., Gaiteri, C., Wennström, M., Kumar, A., Hristovska, I., Spotorno, N., Salvado, G., Strandberg, O., Mathys, H., Tsai, L.-H., et al.: Proteomic changes in alzheimer’s disease associated with progressive a*β* plaque and tau tangle pathologies. Nature neuroscience 27(10), 1880–1891 (2024)

[62] Johnson, E.C., Bian, S., Haque, R.U., Carter, E.K., Watson, C.M., Gordon, B.A., Ping, L., Duong, D.M., Epstein, M.P., McDade, E., et al.: Cerebrospinal fluid proteomics define the natural history of autosomal dominant alzheimer’s disease. Nature medicine 29(8), 1979–1988 (2023)

[63] Guo, Q., Ping, L., Dammer, E.B., Duong, D.M., Yin, L., Xu, K., Shantaraman, A., Fox, E.J., Golde, T.E., Johnson, E.C., et al.: Heparin-enriched plasma proteome is significantly altered in alzheimer’s disease. Molecular neurodegeneration 19(1), 67 (2024)

[64] Sun, L., Wei, G., Ji, F., Ding, Y., Fan, J., Xu, Y., He, C., Zhou, Y., Liu, Z., Sun, Z., et al.: Proteome-wide association study identifies novel alzheimer’s disease-associated proteins. Journal of Alzheimer’s Disease, 13872877251409352 (2024)

